# Hemin Inhibits the Activation of STING in Macrophages by Inducing HO-1, Promoting Endometriosis Development

**DOI:** 10.1101/2025.10.25.683839

**Authors:** QL Mo, RS Chang, LY Zhang, L Huang, W Huang, XY Liang, XL Xue, QB Zhang, XR Hou, YC Lin, ZL Zhou, YW Chen, W Zhu, LW Hao, S Wu

## Abstract

**Introduction:** Endometriosis (EMS) is an estrogen-dependent inflammatory disorder characterized by immune dysregulation. This condition profoundly affects the quality of life and reproductive health of nearly 200 million women of reproductive age worldwide and poses significant clinical challenges due to lack of diagnosis tools and effective treatment. Peritoneal macrophages play a central role in promoting the initiation and progression of EMS. Bioinformatics studies have indicated that the STimulator of INterferon Gene (STING) was downregulated in the macrophages in EMS, and this influenced their polarization and functional. However, the specific contribution of STING pathway alterations to EMS pathogenesis remains unclear.

**Methods:** We integrated scRNA-seq data from EMS patients and healthy controls-including lesions, eutopic endometrium, non-lesional tissue, and peritoneal fluid from EMS patients and healthy controls(n = 19,291 total samples)-to analyze macrophage STING expression and function. Mouse EMS model was generated by injecting uterus fragments into WT or STING^−/−^ mice. Peritoneal fluid was collected and macrophage proportions and subtypes were assessed via flow cytometry. Macrophages were depleted in STING^−/−^ mice prior to EMS induction to evaluate their role in disease progression. In vitro, hemin was used to treat macrophages to investigate how heme oxygenase-1 (HO-1) modulates the STING pathway, assessed through immunoblotting, co-immunoprecipitation, ELISA, and multiplex immunofluorescence.

**Results:** Bioinformatic analysis revealed that macrophages within the EMS lesions are primarily derived from peritoneal macrophages and exhibit decreased STING expression, which correlated negatively with HO-1. STING^−/−^ mice developed more numerous and larger EMS lesions, accompanied by a decreased proportion of large peritoneal macrophages and an increase in small peritoneal macrophages. A marked elevation in M2-type macrophages within the lesions was detected in STING^−/−^ mice. In vitro, hemin-induced HO-1 suppressed STING pathway activation in macrophages. Mechanistically, HO-1 inhibited the translocation of STING from the endoplasmic reticulum to the Golgi, thereby suppressing STING signaling and facilitating EMS.

**Conclusion:** Macrophages are essential for lesion formation in STING^−/−^ mice. In vitro, hemin promotes EMS progression via HO-1 – mediated suppression of the STING pathway. Our findings identify the heme-HO-1-STING axis as a key immunomodulatory pathway in EMS and suggest its potential as a therapeutic target.

Graphical abstract (Created in BioRender.com) Retrograde menstruation leads to the accumulation of heme in the peritoneal cavity, which upregulates HO-1 expression in peritoneal macrophages. HO-1 inhibits the translocation of STING from the endoplasmic reticulum to the Golgi apparatus in macrophages, thereby suppressing the STING pathway and promoting the development and progression of EMS. The progression of EMS, in turn, reinforces this mechanism, creating a self-sustaining vicious cycle.

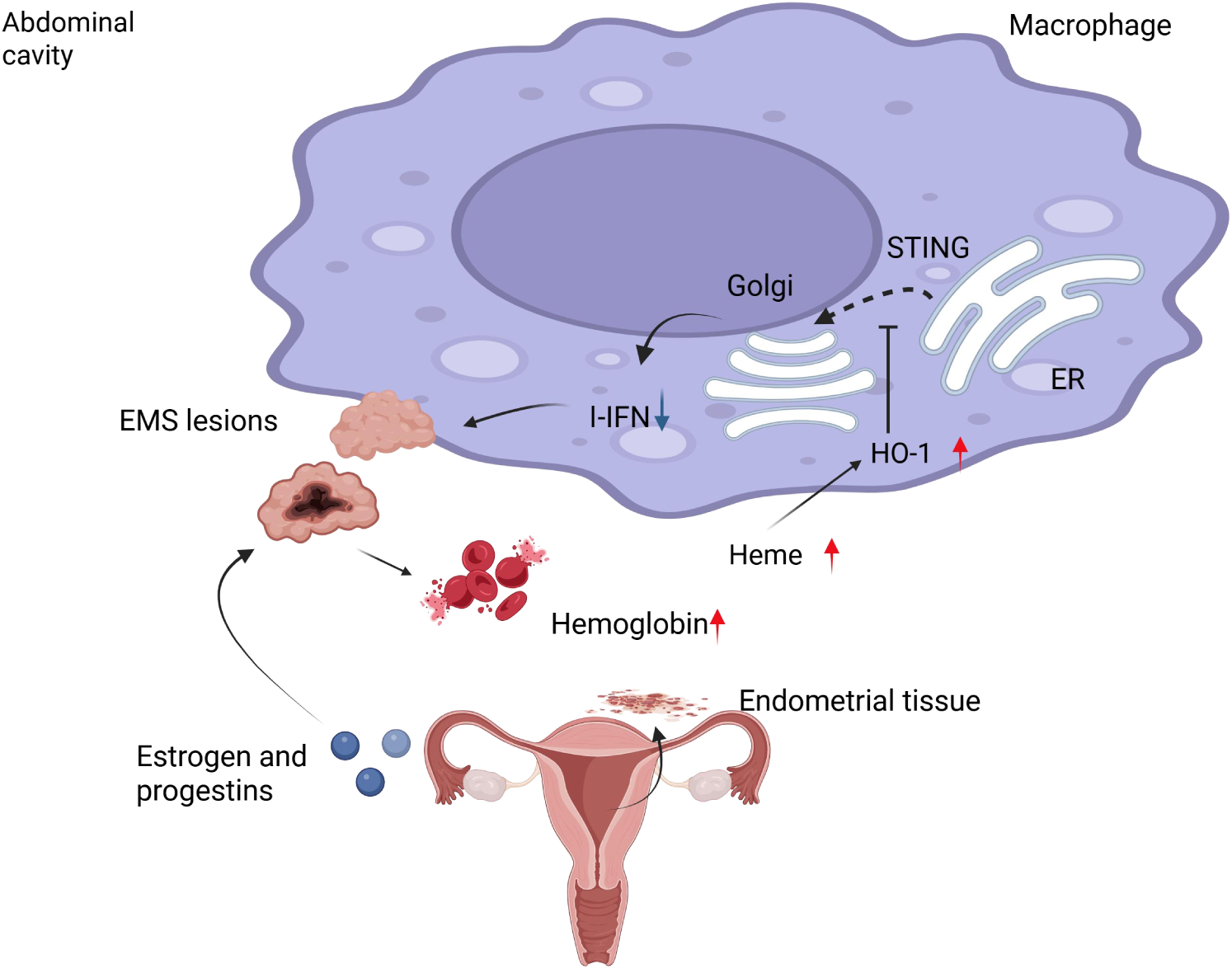

## Introduction

Endometriosis (EMS) is an estrogen-dependent chronic inflammatory gynecological disorder, characterized by the presence of endometrial-like tissue outside the uterine cavity^1^. It affects approximately 5-10% of women of reproductive age, with a recurrence rate of about 30%, often leading to severe dysmenorrhea and significantly impairing patients’ quality of life and fertility^2^. Although benign, EMS exhibits several tumor-like characteristics, such as metastatic behavior, local invasion, proliferative capacity, angiogenesis, and impaired apoptosis^3^. Accumulating evidence indicates that EMS is associated with an increased risk of several malignancies, including endometrioid and clear cell ovarian carcinomas, non-Hodgkin lymphoma, brain tumors, and endocrine cancers^4^. Especially, women diagnosed with EMS have a significantly elevated risk of developing ovarian cancer, with 2.9% of those affected also exhibiting EMS-associated ovarian cancer^5,6^. Therefore, a deeper understanding of the pathogenesis of EMS is crucial for elucidating its link with autoimmunity, mitigating the risk of malignant transformation, and improving early diagnosis and personalized treatment.

Chronic inflammation, fibrosis, and angiogenesis are recognized as central pathological features of EMS^7^. Under normal conditions, peritoneal macrophages-key immune residents in the peritoneal cavity – play essential roles in clearing ectopic endometrial cells via phagocytosis, inflammatory signaling, antigen presentation, cytotoxic T-cell activation, and tissue repair^8–10^. However, in EMS, macrophages are reduced in number and functionally compromised, leading to inadequate clearance of refluxed endometrial debris, immune escape, and eventual lesion establishment^11^. Macrophages are broadly categorized into two polarization states: the classically activated M1 phenotype and the alternatively activated M2 phenotype. M1 macrophages secrete pro-inflammatory cytokines such as IL-12, IL-23, TNF-α, IL-6, and IL-1β, and are critical for initiating inflammation, pathogen elimination, and antigen presentation^12^. In contrast, M2 macrophages produce anti-inflammatory cytokines including IL-4, IL-13, and IL-10, and contribute to immune modulation, tissue remodeling, wound healing, and resolution of inflammation^12^. Macrophage polarization is highly plastic and influenced by the local microenvironment; for example, M1-dominated responses in early inflammation may shift toward an M2 phenotype in later stages^13^. Notably, multiple studies have identified abundant M2 macrophages in EMS lesions and peritoneal fluid, suggesting their active role in disease progression^14–16^. These M2 macrophages support various pathogenic processes such as tissue repair, angiogenesis, neurite outgrowth, and enhanced proliferation and invasion of ectopic cells^15,17^. Therefore, abnormalities in the phenotypes and functions of macrophages can lead to the development of EMS.

STING is an innate immune sensor that recognizes cytosolic nucleic acids derived from pathogens, damaged tissues, or cellular senescence^18^. Upon detecting aberrant DNA, STING activates downstream signaling pathways that induce inflammatory and type Ⅰ interferon responses, playing key roles in antiviral defense, antitumor immunity, and autoinflammatory disorders. Although STING activation enhances macrophage-mediated clearance of ectopic tissues, excessive activation may result in persistent interferon and cytokine production, contributing to tissue damage and chronic inflammation^19,20^. This has been implicated in autoimmune diseases such as systemic lupus erythematosus and rheumatoid arthritis^20^. Interestingly, suppression of STING signaling has been shown to promote M2 macrophage polarization, whereas STING expression favors an M1 phenotype^21–23^. Additionally, STING activation augments macrophage phagocytic capacity by downregulating SIRPα expression^24^. Despite these advances, the mechanisms through which STING modulates macrophage polarization in EMS remain poorly understood.

Ectopic endometrial tissue responds to hormonal changes, particularly estrogen and progesterone fluctuations^25^. Hormonal fluctuations induce a menstrual-like process of hyperplasia, shedding, and irregular bleeding in ectopic endometrial tissue^26,27^. During this process, the lysis of red blood cells leads to the sustained production of heme. Macrophages are key regulators of heme and iron metabolism, capable of uptake hemoglobin and heme via receptors such as CD163 and CD91^28^. Heme oxygenase-1 (HO-1, encoded by HMOX1) is the rate-limiting enzyme in heme degradation, catalyzing the breakdown of heme into carbon monoxide, biliverdin, and ferrous iron^29^. HO-1 exert multiple regulatory functions in the fields of immunity, oxidative stress, inflammation, and cellular metabolism^30^. Elevated HO-1 expression has been consistently observed in peritoneal cavity and endometriotic lesions of EMS patients^31–33^,, which in turn has been linked to fibrosis progression in established EMS lesions^34^. In addition, HO-1 has been shown to promote the polarization of M2 macrophages and the subsequent development of tissue fibrosis in mature EMS lesions^35–38^. However, the precise mechanism by which the heme – HO-1 axis influences macrophage polarization in EMS remains unclear.

In this study, integrative single-cell analysis identified macrophage STING as a key regulator in EMS that is inversely correlated with HO-1. We delineate a pathogenic “heme-HO-1-STING” axis: heme accumulation upregulates HO-1, which suppresses the STING – IFN-I pathway to drive M2 polarization and disease progression. By combining clinical analysis, multi-omics, and experimental models, our study establishes the therapeutic potential of targeting this axis, redefining STING’s role in EMS and paving the way for novel immunotherapies.

## Results

### 1 Single-cell sequencing analysis revealed that STING was downregulated in ectopic endometrial lesions

STING is critically implicated in chronic inflammation and autoimmune processes. In orther to elucidate whether STING acts as a key regulatory gene in EMS, we performed an integrated reanalysis of single-cell RNA sequencing data from macrophages obtained from EMS patients and healthy volunteers. Macrophages were isolated from the following sources: EMS lesions (n=5783), eutopic endometrium (n=2369), non-lesional abdominal tissue (i.e., tissue not diagnosed as EMS lesions, n=1562), and peritoneal fluid (n=5009) of EMS patients, as well as peritoneal fluid from healthy volunteers (control, n=4568). These cells were pooled and reclassified into two major subsets – endometrial and peritoneal macrophages-comprising 15 distinct clusters (Figure 1A-D).

**Figure 1.**
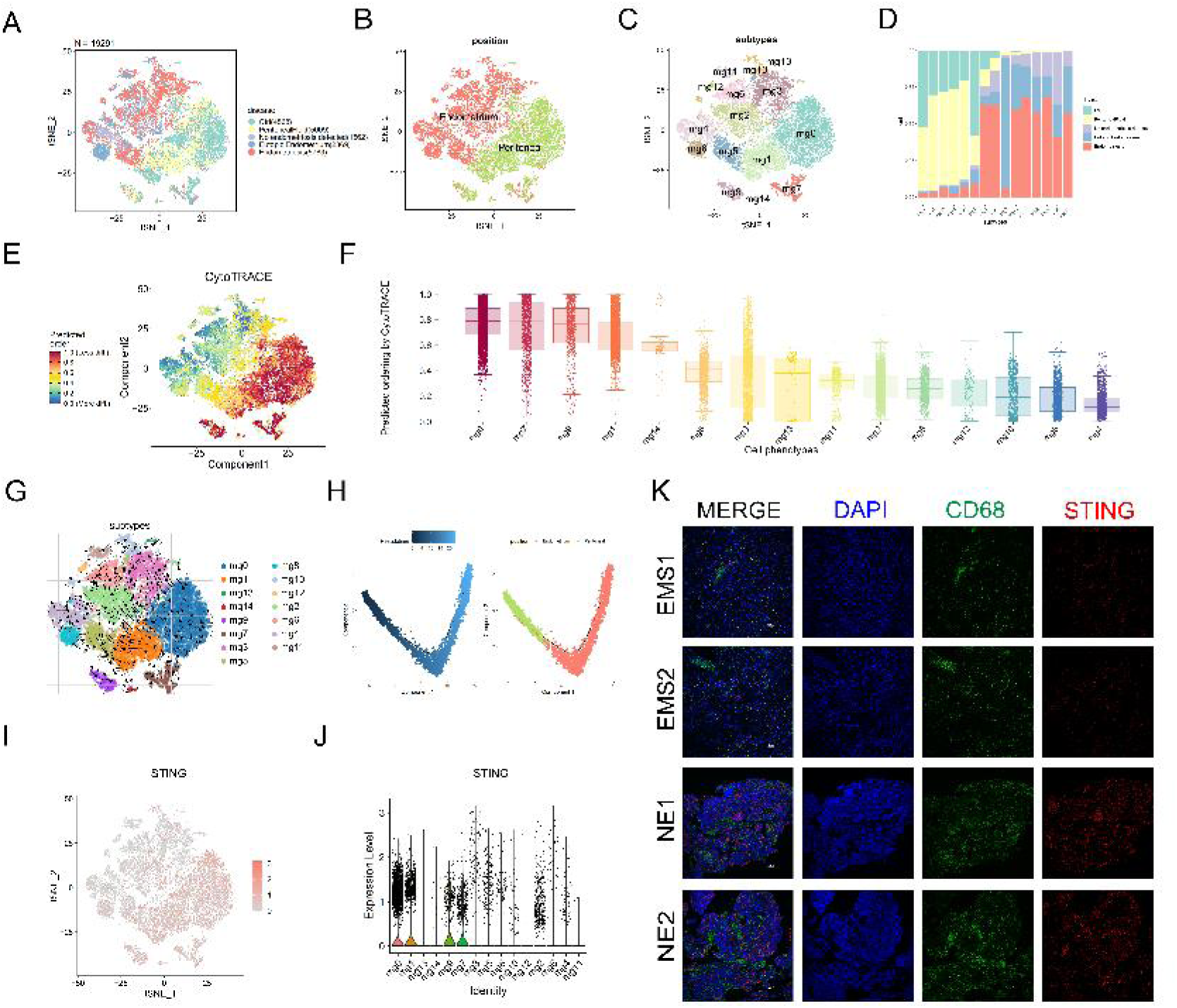
Single-cell sequencing analysis revealed that STING was downregulated in ectopic endometrial lesions. (A–B) UMAP visualization of macrophages from 5 tissue regions and 2 locations. (C) Classification of macrophages into 15 distinct subtypes, shown via UMAP projection. (D) Proportional distribution of the 15 macrophage subtypes across 5 regions. (E–F) Differentiation index scores for the 15 macrophage subtypes. (G) Differentiation trajectories inferred across the 15 macrophage subtypes. (H) Pseudotime trajectory analysis revealing the development of peritoeal macrophages and endometrial macrophages. (I) Spatial distribution of STING expression among macrophage subtypes. (J) Expression levels of STING across the 15 macrophage clusters. (K) Representative mIHC analysis of STING (red) and macrophage (CD68⁺, green) in ectopic lesions from healthy controls and EMS patients. Nuclei are counterstained with DAPI (blue). Scale bar = 100 μm. NE: normal endometrium; EMS: endometriosis.

The differentiation index was lower in the peritoneal macrophage subsets (clusters 0, 1, 7, 9, 14) and higher in the endometrial subsets (clusters 2, 3, 4, 5, 6, 8, 10, 11, 12, 13) (Figure 1E-F). This pattern was further supported by differentiation trajectory analysis (Figure 1G-H), suggesting that macrophages within EMS lesions may originate from peritoneal macrophages.

Subsequently, we assessed the distribution and expression of STING across macrophage subsets (Figure 1I-J). STING expression was found to be lower in endometrial-derived macrophages compared to those derived from the peritoneal cavity. Moreover, multicolor immunohistochemistry of clinical samples confirmed that STING expression was downregulated in EMS lesions relative to normal endometrial tissue (Figure 1K). Based on these findings, we propose that STING expression decreases as macrophages migrate from the peritoneal cavity into the lesions during disease progression, indicating that STING may serve as a pivotal gene in the pathogenesis of EMS.

### 2. STING deficiency promotes the development of EMS

To investigate the role of STING in the pathogenesis of EMS, we established an EMS model using STING^−/−^ and wild-type (WT) mice as recipients, with WT mice serving as donors (Figure 2A). Lesions in STING^−/−^ recipients showed pronounced enlargement compared to those in WT recipients (Figures 2B-C), suggesting that STING deficiency facilitates both the initiation and progression of EMS. Furthermore, lesions in STING^−/−^ mice exhibited more typical pathological features of ectopic endometrium (Figure 2D).

**Figure 2.**
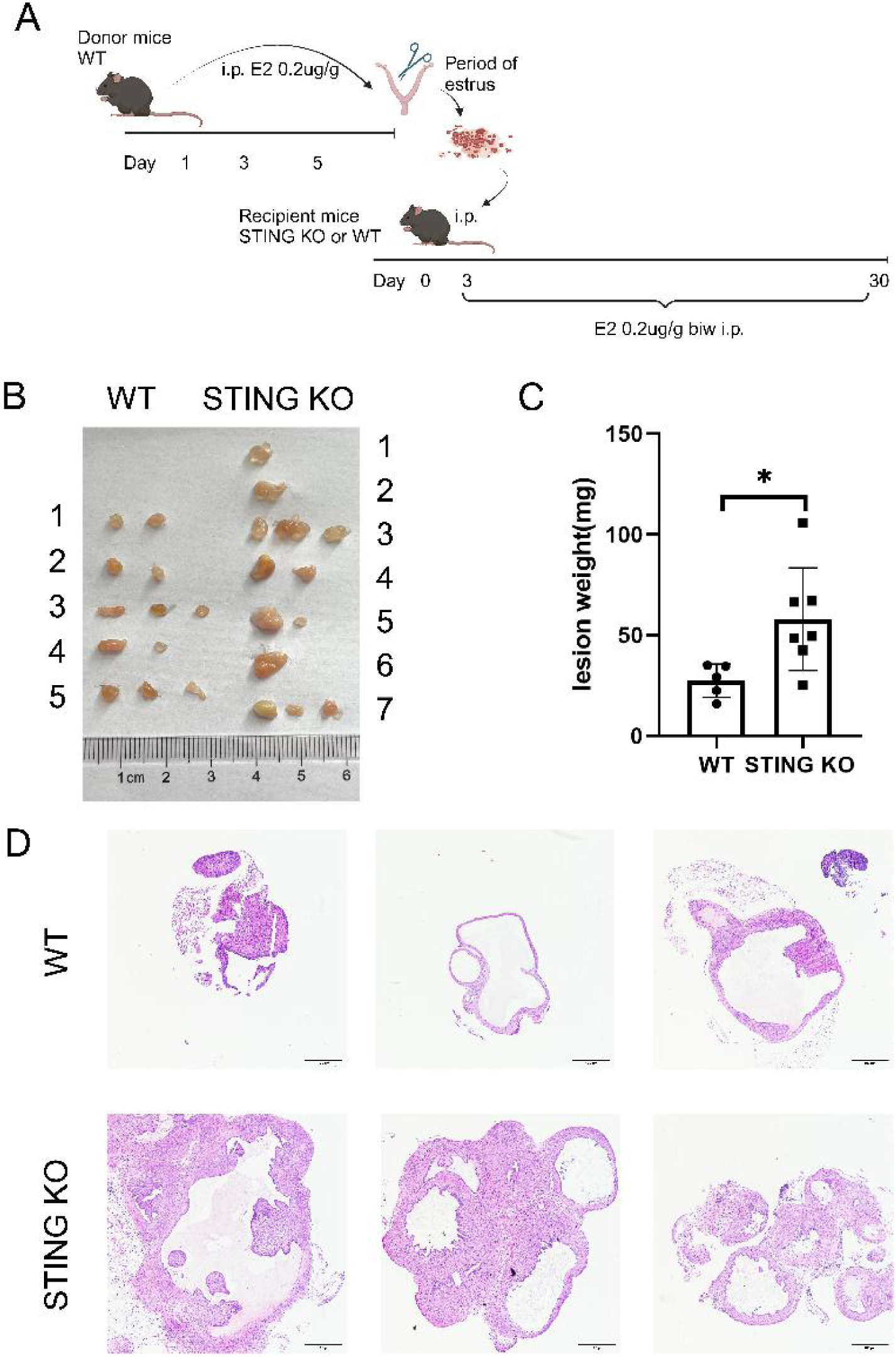
STING deficiency promotes the development of EMS. (A) Schematic diagram of the experimental EMS model in mice. (B) Macroscopic appearance of ectopic lesions one month after induction. (C) Quantification of lesion weights in WT and STING^−/−^ mice. (D) H&E staining of endometrial lesions. Scale bar=100 μm. *p<0.05.

To further assess the influence of donor uterine tissue on EMS development, we employed STING^−/−^ mice as donors in WT and STING^−/−^ recipient mice. When STING^−/−^ donors were used, WT recipients developed larger lesions compared to STING^−/−^ recipients (Supplementary Figure 1A). Moreover, the proportion of peritoneal macrophages remained unaltered across these groups (Supplementary Figure 1C-D). In contrast, lesion size and weight in STING^−/−^ recipients did not differ significantly regardless of whether donors were STING^−/−^ or WT mice (Supplementary Figure 1B).

These results indicate that donor genetic background has limited influence on EMS development in a STING-deficient microenvironment. There is a complex interaction between donor endometrial cells and recipient macrophages in modulating EMS development. Overall, our findings demonstrate that STING deficiency promotes the occurrence and progression of EMS.

### 3. STING deficiency in macrophages promotes the development of EMS

To investigate whether STING in macrophages contributes to the initiation and progression of EMS, a series of experiments were performed. Macrophages were first depleted in STING^⁻/⁻^ mice, followed by the induction of EMS models (Figure 3A). The absence of macrophages in STING^⁻/⁻^ mice led to a significant reduction in lesion formation (Figure 3B), indicating that macrophages play a crucial role in EMS development in these mice.

**Figure 3.**
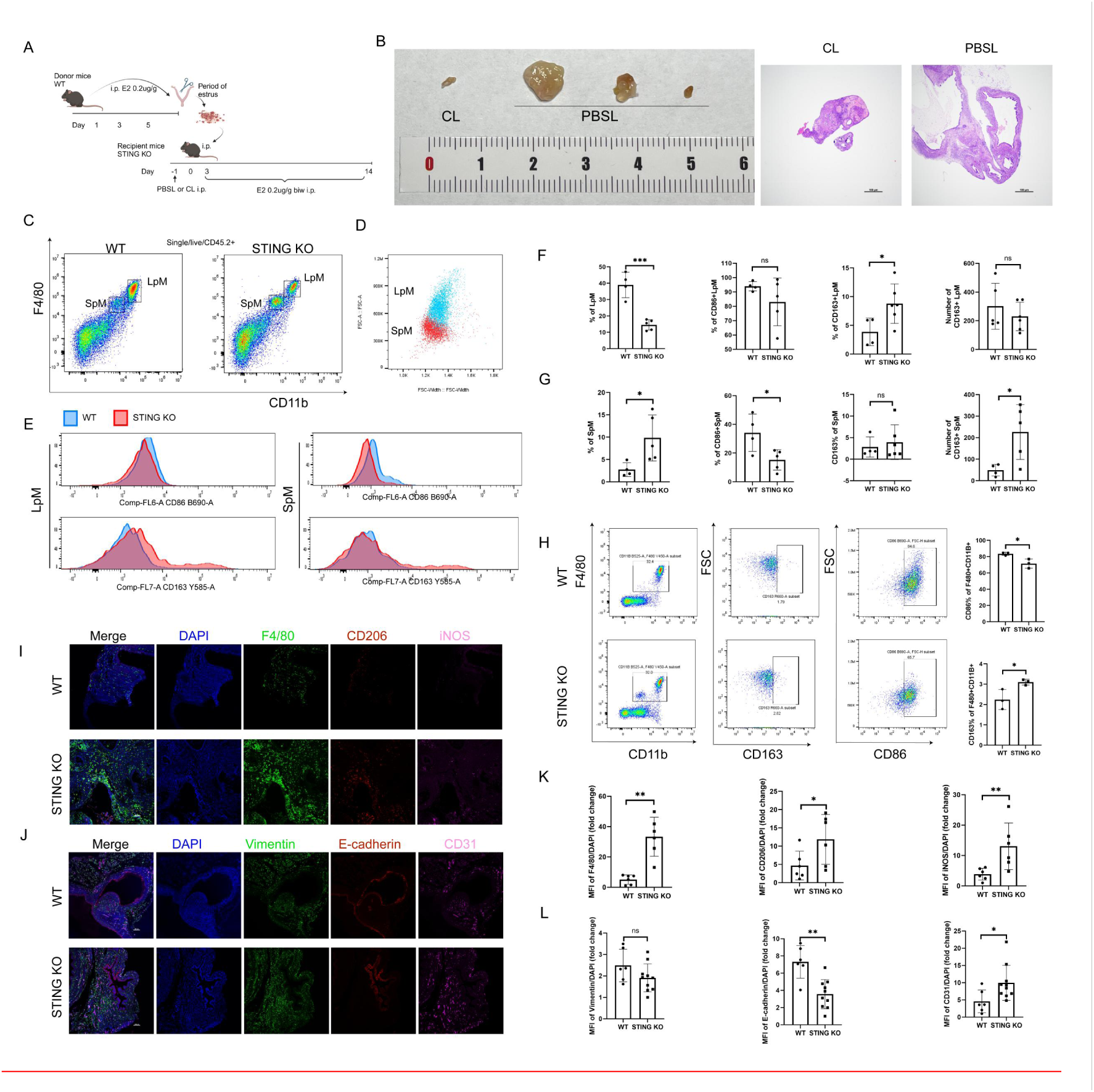
STING deficiency in macrophages promotes the development of EMS. (A) Schematic diagram of the experimental EMS model in STING^−/−^ mice. (B) Macroscopic appearance and H&E staining of lesions from STING^−/−^ mice on day 14 (CL group, n = 3; PBSL group, n = 3). (C) Gating strategy for identifying small peritoneal macrophages (SpM) and large peritoneal macrophages (LpM) (singlets/live/CD45^+^/F4/80^+^/CD11b^+^) in peritoneal fluid from EMS-induced mice. (D) Cell size comparison between LpM and SpM. (E) Flow cytometry plots showing CD86⁺ and CD163⁺ expression within LpM (left) and SpM (right). (F) Relative abundance of LpM populations in peritoneal fluid, quantification of CD86^+^ or CD163^+^ LpM. (G) Relative abundance of SpM populations in peritoneal fluid, quantification of CD86^+^ or CD163^+^ SpM. (H) Flow cytometry plots showing CD86^+^ or CD163^+^ expression in peritoneal macrophages from WT or STING^−/−^ mice. (I) Representative mIHC of lesions showing DAPI (blue), F4/80 (green), CD206 (red), and iNOS (magenta). Scale bar=100 μm. (J) Representative mIHC of lesions showing DAPI (blue), vimentin (green), E-cadherin (red), and CD31 (magenta). Scale bar=100 μm. (K) Quantification of fluorescence intensity for F4/80, iNOS, and CD206 in lesions. (L) Quantification of fluorescence intensity for vimentin, E-cadherin, and CD31 in lesions. LpM, large peritoneal macrophage; SpM, small peritoneal macrophage; CL, clodronate liposomes; PBSL, PBS liposomes. *p < 0.05, ** p < 0.01, *** p < 0.001, ns: not significant.

We next analyzed the proportions and specific markers of macrophages in the peritoneal fluid of STING^⁻/⁻^ and WT mice with EMS. Two distinct subsets of peritoneal macrophages were identified. Based on macrophage marker expression and cell size, the F4/80ˡᵒʷ CD11bˡᵒʷ population was classified as small peritoneal macrophages (SpM), which were increased in STING^⁻/⁻^ mice. In contrast, the F4/80ʰⁱᵍʰ CD11bʰⁱᵍʰ population was classified as large peritoneal macrophages (LpM), which were decreased in STING^⁻/⁻^ mice (Figure 3C-D).

Compared to WT mice, STING^⁻/⁻^ mice exhibited a significantly higher proportion of CD163⁺ macrophages within the LpM subset (Figure 3E-F). Conversely, in the SpM subset, the proportion of CD86⁺ macrophages was significantly reduced, while that of CD163⁺ macrophages was increased in STING^⁻/⁻^ mice (Figure 3E, G). Notably, SpM were present in untreated STING^⁻/⁻^ mice but not detected in WT mice, with elevated CD163^+^ and reduced CD86^+^ proportions under basal conditions (Figure 2H).

Moreover, an increased infiltration of macrophages was observed in the lesions of STING^⁻/⁻^ mice (Figure 3I), with a more pronounced increase in M2 macrophages (marked by CD206) than in M1 macrophages (marked by iNOS). Additionally, CD31 fluorescence was significantly elevated, while E-cadherin fluorescence was reduced in STING^⁻/⁻^ mice (Figure 3J), suggesting enhanced angiogenesis and epithelial–mesenchymal transition (EMT) within EMS lesions. Co-localization of M2 macrophages with CD31 was also detected in the lesions of STING^⁻/⁻^ mice (Supplementary Figure 2). Further analysis of macrophage subsets in lesions and spleens of EMS mice consistently showed higher lesion cell viability and a decreased M1/M2 ratio in STING^⁻/⁻^ mice (Supplementary Figure 3).

In summary, these results demonstrate that STING deficiency in macrophages reprograms macrophage polarization, promotes angiogenesis and EMT, and facilitates the development of EMS.

### 4 Negative correlation between HO-1 and STING expression in EMS associated macrophages

To investigate the potential correlation between HO-1 and STING expression in EMS-associated macrophages, the distribution and expression of HO-1 across macrophage subsets were detected. Notably, although both STING and HO-1 were generally expressed at higher levels in peritoneal-derived macrophages compared to endometrial-derived macrophages, we detected a significant inverse correlation between HO-1 expression and levels of STING or ISG15 specifically within peritoneal macrophages (Figure 4A-C).

**Figure 4.**
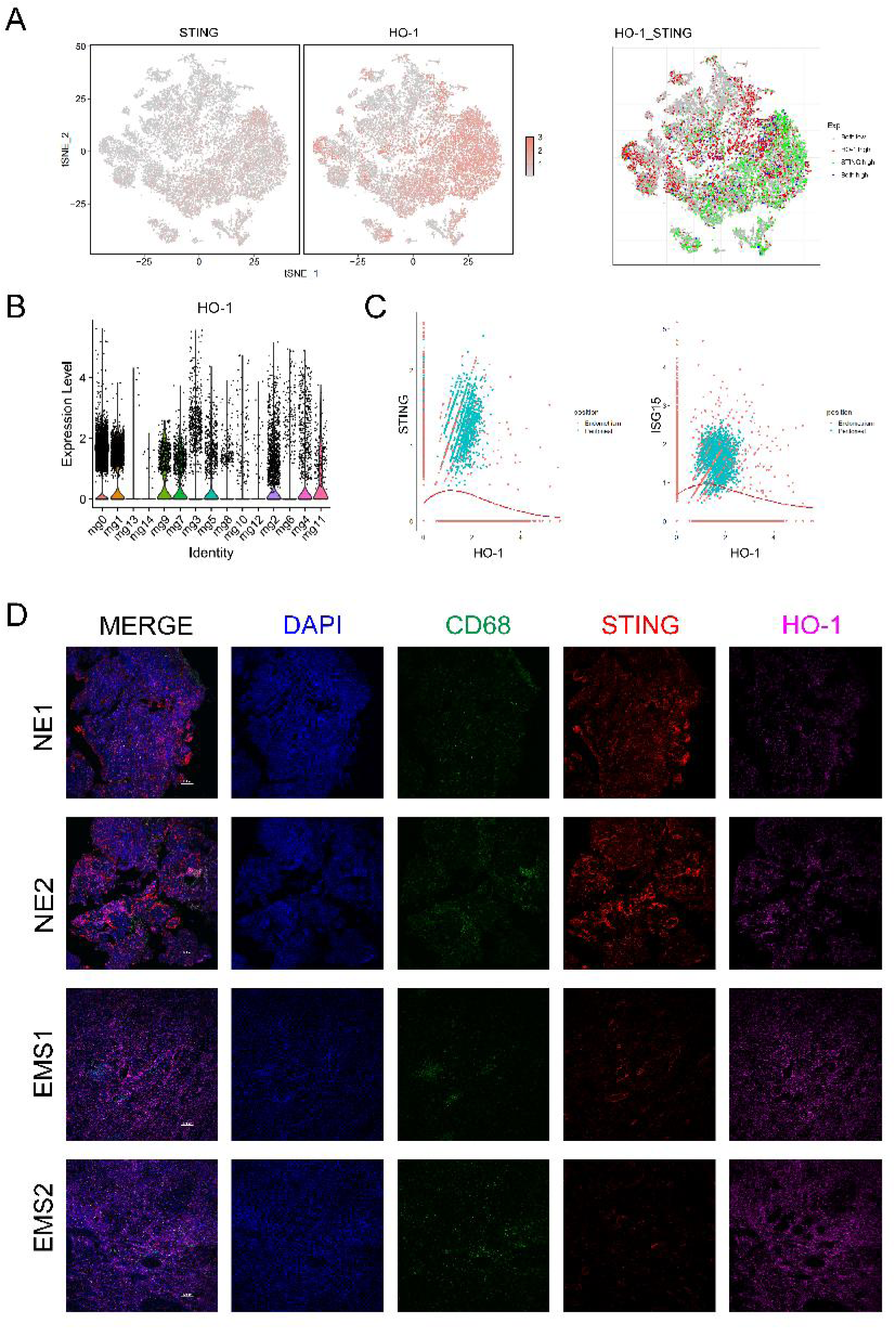
Negative correlation between HO-1 and STING expression in EMS associated macrophages. (A) Spatial distribution of STING and HO-1 expression among macrophage subtypes. (B) Expression levels of HO-1 across the 15 macrophage clusters. (C) Correlation analysis between HO-1 expression and STING or ISG15 expression in peritoneal and endometrial macrophages. (D) Representative mIHC analysis of STING (red), HO-1(magenta), and macrophage (CD68⁺, green) co-localization in ectopic lesions from healthy controls and EMS patients. Nuclei are counterstained with DAPI (blue). Scale bar = 100 μm. NE: normal endometrium; EMS: endometriosis.

Furthermore, ectopic lesions exhibited upregulated HO-1 expression and downregulated STING expression compared to normal endometrium (Figure 4D). Collectively, these results indicate a negative correlation between HO-1 and STING expression in macrophages derived from EMS patients.

### 5. High-concentration hemin induces HO-1 expression and inhibits STING-IFN-Ⅰ pathway activation

We hypothesized that heme induces HO-1 expression and modulates the STING-IFN-Ⅰ pathway. To test this, peritoneal macrophages and Raw264.7 cells were treated with varying concentrations of hemin. Initial assays ruled out potential effects of high hemin concentrations on cell proliferation and cytotoxicity (Supplementary Figure 4). In Raw264.7 cells, HO-1 expression was induced in a concentration-dependent manner. Notably, high-concentration hemin (50 μM) treatment significantly suppressed protein levels of the STING signaling pathway (Figure 5A-B). Similarly, hemin pre-treatment profoundly inhibited STING pathway activation in peritoneal macrophages that had been stimulated with STING agonists (Figure 5C). Furthermore, hemin pre-treatment in both peritoneal macrophages and bone marrow-derived macrophages (BMDMs) markedly reduced IFN-β production induced by STING agonists (Figure 5D). Subsequent flow cytometry analysis revealed that STING agonist treatment in BMDMs enhanced both the proportion and fluorescence intensity of CD86⁺cells, indicating promoted M1 macrophage polarization. This effect was attenuated by hemin pre-treatment (Figure 5F). Immunofluorescence staining also demonstrated co-localization of HO-1 and STING proteins within cells (Figure 5E).

**Figure 5.**
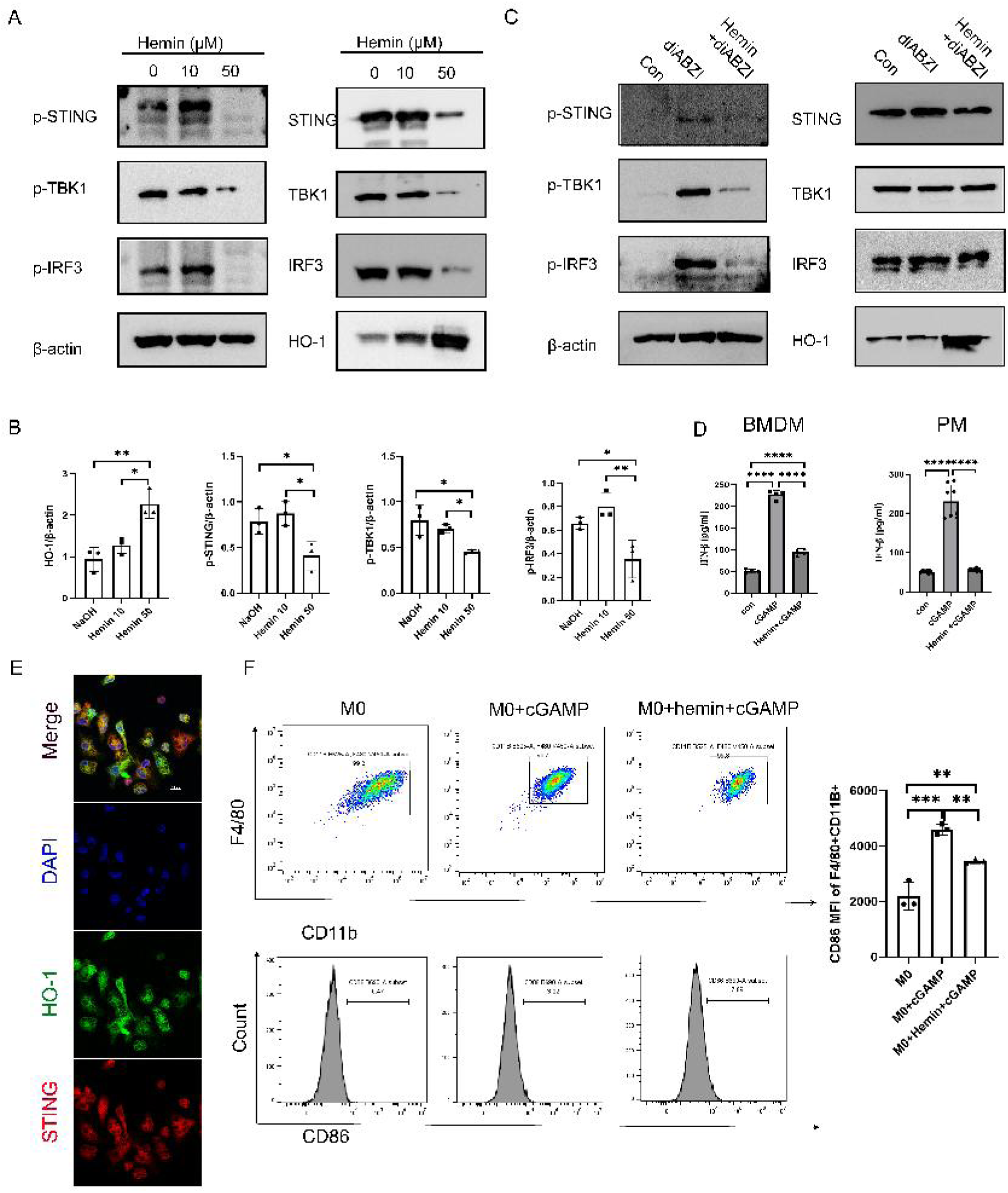
High-concentration hemin induces HO-1 expression and inhibits STING-IFN-Ⅰ pathway activation. (A–B) Immunoblot analysis of HO-1 and STING pathway proteins in Raw264.7 cells treated with increasing concentrations of hemin. (C) Immunoblot analysis of STING signaling proteins in PMs under the indicated treatments. (D) IFN-β secretion measured by ELISA in the supernatant of BMDMs and PMs after various treatments. (E) Representative confocal microscopy images showing subcellular localization of STING (red) and HO-1 (green) in peritoneal macrophages. Nuclei are stained with DAPI (blue). (F) Flow cytometry analysis of CD86⁺ macrophages. Gating strategy: single cells → live → CD45⁺ → F4/80⁺ → CD11b⁺. Right panel: quantitative analysis of CD86 fluorescence intensity in BMDMs under different treatments. Data are representative of at least three independent experiments. *p < 0.05, **p < 0.01, ****p < 0.0001. BMDMs: bone marrow-derived macrophage; PMs: peritoneal macrophage.

In summary, hemin treatment not only upregulates HO-1 expression but also inhibits STING-IFN-Ⅰ pathway activation and restricts M1 macrophage polarization. Additionally, HO-1 and STING were observed to be co-localized. However, further studies are needed to determine whether hemin suppresses STING-IFN-Ⅰ signaling specifically through HO-1 induction.

### 6. HO-1 inhibits STING pathway activation by impairing ER-to-Golgi translocation of STING

To further investigate whether hemin inhibits STING pathway activation through the induction of HO-1 expression, we performed co-immunoprecipitation assays to examine the interaction between HO-1 and STING. As shown in Figure 6A, HO-1 was found to interact with STING in BMDMs. This interaction was reduced upon stimulation with STING agonists but was significantly enhanced by pre-treatment with hemin (Figure 6B). Consistent with this, immunofluorescence analysis revealed that STING agonist stimulation reduced the co-localization of HO-1 and STING, whereas hemin pre-treatment restored their co-localization even in the presence of STING agonists (Figure 6C).

**Figure 6.**
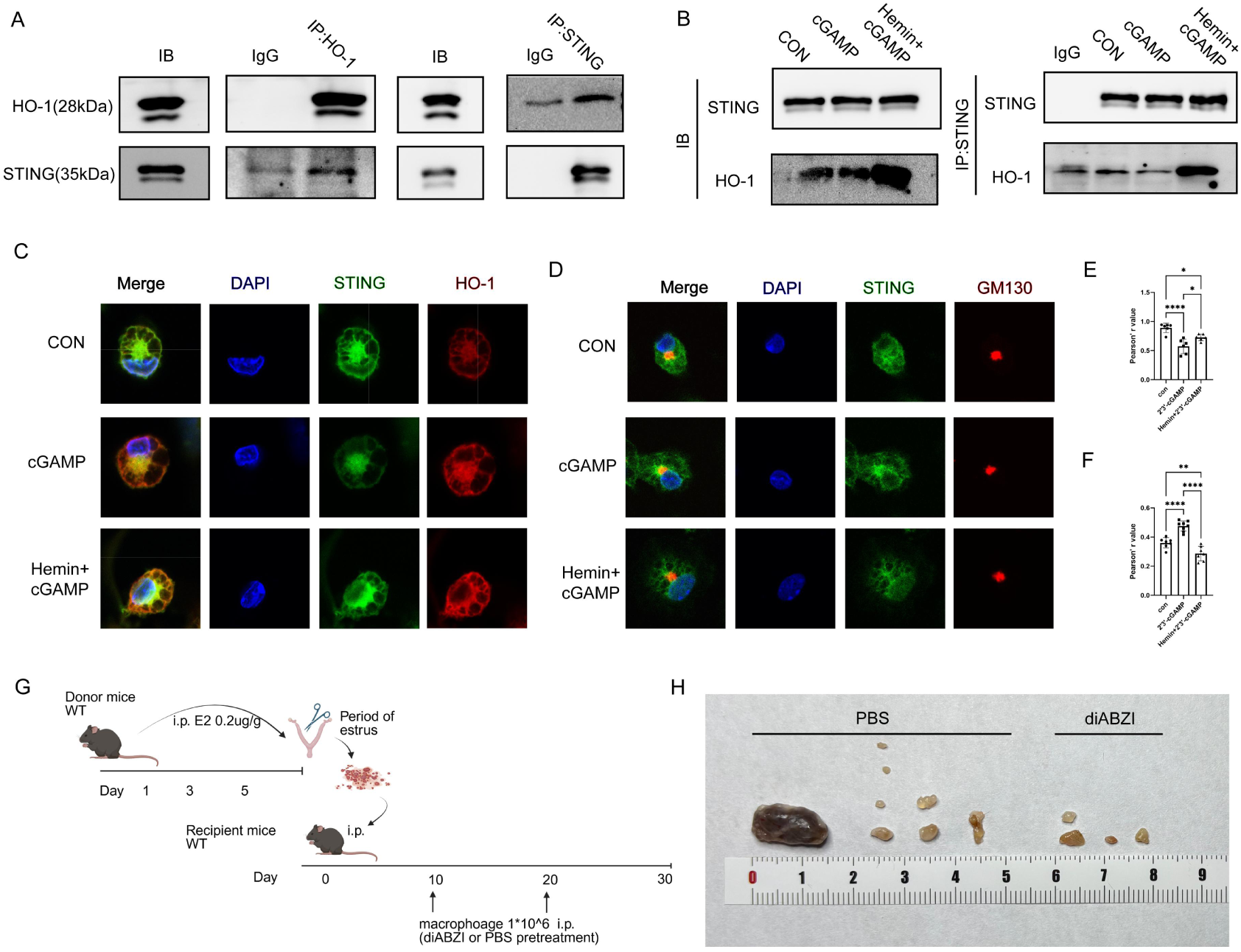
HO-1 inhibits STING pathway activation by impairing ER-to-Golgi translocation of STING. (A) Interaction between endogenous STING and HO-1 in unstimulated BMDMs was assessed by co-immunoprecipitation. (B) Co-immunoprecipitation analysis of the STING – HO-1 interaction in BMDMs treated with 2’3’-cGAMP or hemin and 2’3’-cGAMP. (C) Representative confocal microscopy images showing subcellular localization and co-localization of STING (green) and HO-1 (red) in peritoneal macrophages under the indicated treatments. (D) Confocal microscopy images illustrating STING (green) and Golgi marker GM130 (red) co-localization in peritoneal macrophages following various treatments. (E) Quantification of cells exhibiting STING-HO-1 co-localization. Data are presented as mean ± SEM; *p < 0.05, **p < 0.01, **p < 0.0001. (F) Statistical analysis of cells for STING-GM130 co-localization. Values represent mean ± SEM; *p < 0.05, **p < 0.01, **p < 0.0001. (G) Schematic diagram of the experimental protocol for establishing the EMS model in WT mice and administering the indicated treatments. (H) Representative macroscopic images of peritoneal lesions from WT mice with or without the indicated treatments (PBS group, n = 4; diABZI group, n = 4).

Subsequent immunofluorescence analysis also showed that hemin pre-treatment reduced the co-localization between GM130 (a Golgi marker) and STING following agonist stimulation, suggesting that hemin impairs the translocation of STING from the endoplasmic reticulum (ER) to the Golgi, thereby inhibiting downstream STING pathway activation (Figure 6D).

In summary, these results demonstrate that hemin inhibits STING pathway activation and type I IFN production by upregulating HO-1 expression. HO-1 interacts with STING and inhibits its translocation from the ER to the Golgi, thereby suppressing downstream signaling.

### 7. STING-activated macrophages alleviate the development of EMS in mice

The therapeutic potential of STING activation in macrophages for mitigating EMS progression was subsequently investigated. Peritoneal macrophages were isolated from WT mice, followed by purification via flow cytometry (Supplementary Figure 5). After a 2-hour in vitro pre-treatment with either the STING agonist diABZI or PBS, these macrophages were intraperitoneally transferred into WT mice with induced EMS (Figure 6G). Administration of diABZI-activated macrophages resulted in a significant decrease in the number and volume of EMS lesions (Figure 6H). Taken together, these results demonstrate that STING activation in macrophages can suppress the development of endometriotic lesions in a mouse model.

## Discussion

In this study, we demonstrate that STING deficiency in macrophages promotes the development of EMS in vivo. In vitro, heme induces HO-1 expression, which inhibits the translocation of STING from the ER to the Golgi, thereby suppressing the STING-IFN-I pathway. Furthermore, treatment with STING agonists reduced lesion formation in a murine EMS model. Our findings reveal a novel link between heme metabolism and STING signaling in EMS and uncover an unexpected anti-inflammatory role of STING suppression in EMS-associated macrophages.

Immune dysregulation is one of the hallmarks of EMS. The disease triggers local inflammation and cytokine secretion, recruiting immune cells to clear endometrial lesions^39^. STING is a key immune regulator influencing macrophage differentiation and phagocytosis^40,41^. Although STING pathway inhibition has been shown to ameliorate inflammatory and autoimmune disorders^19,20^, we found that loss of STING in macrophages exacerbated EMS. Notably, STING^−/−^ mice exhibited a significant increase in the SpM subset, which is monocyte-derived and typically exhibits an M2-like phenotype. In contrast, the LpM subset-embryonically derived and often M1-like-was reduced. These observations align with previous reports showing an increased SpM/LpM ratio in EMS mouse models^42^. Other studies support that STING inhibition promotes M2 polarization, whereas STING activation drives M1 polarization^21–23^. For example, Su et al. reported that STING inhibitors suppressed M1 polarization while enhancing M2 differentiation in hepatic macrophage^43^. Moreover, STING activation enhances phagocytic capacity^24,44^, suggesting that intact STING signaling may support pathogen clearance and exert protective roles in EMS.

Previous studies present seemingly conflicting roles for STING in EMS. One report indicated that STING is upregulated in human and rat ectopic endometrium, promoting autophagy, migration, and invasion in human endometrial stromal cells (HESCs)^45^. Another study found that low STING expression facilitated HESC invasion and migration via the STING/IRF3/IFN-β1 pathway in eutopic endometrium from EMS patients^46^. These discrepancies may stem from cell-type-specific functions: in stromal cells, STING modulates local inflammation, tissue repair, and fibrosi, whereas in immune cells, it regulates systemic immunity and pathogen clearance^47^. Additionally, STING may exert context-dependent effects influenced by the cellular microenvironment, disease stage, or source of DNA damage^48^. Furthermore, our results suggest that STING agonists may have repurposing potential for EMS treatment. However, systemic STING activation carries risks; thus, local administration strategies and optimal treatment timing warrant further investigation.

We further demonstrated that high concentrations of heme induce HO-1 expression, which inhibits the STING-IFN-I pathway. HO-1 plays pleiotropic roles in immunity, oxidative stress, inflammation, and cellular metabolism^30^. A previous CRISPR screen identified HO-1 as a negative regulator of IFN-I production^49^, ER-anchored HO-1 disrupts STING polymerization and trafficking to the Golgi.^49^, supporting the role of HO-1 in suppressing STING-dependent IFN responses. Notably, combining STING agonists with HO-1 inhibitors synergistically enhances antitumor immunity^50^. However, one study reported that heme can bind STING directly and activate STING–IFN-β signaling in cerebral endothelial cells^51^. These opposing effects may be due to differences in heme concentration, exposure time, or cell type, suggesting that heme may modulate STING through multiple mechanisms. The dual role of heme in regulating STING implies that therapeutic strategies should be tailored based on the inflammatory context.

This study has several limitations. First, the lack of longitudinal human data prevents establishing a causal relationship between heme levels and STING activity in EMS. Second, the dynamics of the heme-HO-1-STING axis across different disease stages remain unclear. Third, the influence of heme-STING interactions in other peritoneal cells (e.g., endometrial stromal cells) requires further exploration. Finally, the origin of macrophages within EMS lesions was not traced in this study, though prior work suggests contributions from endometrial tissue, LpM, and SpM^52^.

From a clinical perspective, EMS remains challenging due to diagnostic delays and high recurrence rates^53^.A deeper understanding of its pathogenesis is essential. In this study, we integrated single-cell RNA sequencing of macrophages from EMS patients with mechanistic studies in heme-treated macrophages to: (i) delineate the role of STING in EMS-associated macrophages; (ii) demonstrate that heme inhibits STING activation by blocking ER-to-Golgi trafficking; and (iii) evaluate the therapeutic potential of STING agonists in a mouse EMS model. Our findings position STING as a double-edged molecule in EMS and may inform future STING-targeted immunotherapies.

## Conclusion

STING deficiency in macrophages promotes EMS progression. Mechanistically, heme-induced HO-1 interacts with STING, inhibiting its activation and downstream signaling.

## Methods

### 1. Data acquisition and processing

The single-cell mRNA sequence (scRNA-seq) dataset of peritoneal fluid of endometriosis was free downloaded at NCBI’s Gene Expression Omnibus, with the Accession Number of PRJNA713993. The scRNA-seq dataset and Spatial mRNA sequence (stRNA-seq) dataset of endometriosis was free downloaded at GEO database, with the Accession Number of GSE213216. Both scRNA-seq datasets were processed with the Seurat package (version 5.0.3). Macrophages were extracted via co-expression of PTPRC/CD45, CD68, CD14, LYZ, CLEC10A, C1QA and FCGR3A/CD16 and integrated with the “Harmony” algorithm in R software (version 4.4.0)^54,55^. Regarding the integrated macrophage data, cell profiles were filtered employing the criteria of the original publications^54,56,57^. Finally, 9577 macrophages from peritoneal fluid of endometriosis and 9714 from endometriosis were obtained. The curated macrophages were further profiled from control(n=4568), peritoneal fluid(5009), no endometriosis detected (n=1562), eutopic endometrium (n=2369) and endometriosis (n=5783) according to the original publications^56,57^.

### 2. scRNA-seq analysis

The Seurat (version 5.1.0)^58^ was employed to perform single-cell analysis. Macrophages were divided into 15 clusters using the “FindNeighbors” and “FindClusters” functions, and a resolution of 0.3 was set to increase accuracy. Then, uniform manifold approximation and projection (UMAP) was used to represent the high-dimensional cell data in a two-dimensional form, which groups together cells with similar expression patterns and displays cells with different patterns separately. To annotate macrophage subclusters, marker genes specific to each cell type were determined through the FindAllMarkers function, configured with the following parameters: logfc.threshold = 0.5, only.pos = TRUE, and min.pct = 0.25. The proportion of cells in each macrophage cluster from the peritoneal fluid and endometrium, as well as their presence in the control group, peritoneal fluid, no endometriosis detected, eulopic endometrium and endometriosis, was used to identify macrophages that specifically migrate from peritoneal fluid to the uterus during the occurrence of endometriosis.

### 3. Differentiation potential analysis

The CytoTRACE algorithm, developed by Gulati et al.^59^, is an advanced tool for analyzing scRNA-Seq data. Its core function is to capture, refine, and quantify gene expression levels highly correlated with single-cell gene counts. After the CytoTRACE computation, each macrophage is assigned a score that describes its stemness state. As a reliable computational method, CytoTRACE accurately predicts cell differentiation states and has been validated in large-scale datasets, outperforming traditional stemness assessment algorithms. In this study, we used the R package CytoTRACE2 v1.1.0 to calculate the CytoTRACE scores, with scores ranging from 0 to 1. A higher score indicates stronger stemness (lower differentiation), while a lower score suggests weaker stemness (higher differentiation).

To explore complex cellular dynamics in response to environmental changes, CellRank^60^ was utilized via the CytoTRACE kernel to compute transition matrices based on CytoTRACE scores. The CytoTRACE kernel generates a transition matrix that reflects the likelihood of cells transitioning from one state to another.This integration allows for a more comprehensive analysis of cellular trajectories by estimating the probabilities of transitioning between different cell states. The transition matrix can be visualized in a low-dimensional embedding, providing insights into the differentiation pathways and relationships between different cell types.

### 4. Cell pseudotime analysis

The R package Monocle v2.32.0 ^61^ was used to perform cellular pseudotime and trajectory analyses, investigating the differentiation pathways of various annotated cell subgroups, as well as the genes associated with different trajectory patterns. The “DDRTree” function was applied to infer pseudotime trajectories between different cell subtypes and to generate trajectory maps depicting potential developmental relationships among these cell populations.

### 5. Animals

Female C57BL/6JGpt mice aged 6-8 weeks were purchased from GuangDong GemPharmatech Co., Ltd. All animals were maintained under specific pathogen-free conditions at a temperature of 20-26 °C with a 12-hour light/dark cycle. Food and water were provided ad libitum.

### 6. Induction of endometriosis

Six-to eight-week-old female C57BL/6JGpt or STING^−/−^ donor mice received intraperitoneal injections of estrogen (0.2 µg/g body weight in 200 µl PBS; Cayman, 10006315) on days 1, 3, and 5. Donor mice in estrus were euthanized, and their uteri were excised and minced into fragments measuring 1-3 mm. Uterine fragments from on STING^−/−^ donor mouse were intraperitoneally inoculated into each recipient (C57BL/6JGpt or STING^−/−^ mice) in 500 µl PBS. From day 4 onward, recipient mice received intraperitoneal injections of estrogen (0.2 µg/g in 200µl PBS) every three days. One month after inoculation, mice were euthanized for analysis.

### 7. Macrophage depletion in STING−/−mice

Prior to endometriosis induction, 6-8 week-old STING^−/−^ female mice were injected intraperitoneally with either clodronate liposomes (CL) or PBS liposomes (PBSL) (80 µl per mouse; FormuMax, F70101C-NC). On the following day, these mice were used to establish endometriosis models. Two weeks later, the mice were euthanized and analyzed.

### 8. Isolation and culture of mouse peritoneal macrophages

Peritoneal macrophages were isolated from 6-8 week-old female mice via peritoneal lavage as previously described^62^. Cells were filtered through a 70-µm strainer, centrifuged at 600g for 5 minutes, and resuspended in RPMI 1640 medium (Gibco) supplemented with 10% FBS, 1% penicillin, and 1% streptomycin. Cells were cultured at 37°C in a 5% CO₂ incubator.

### 9. Supplementation of diABZI-stimulated macrophages in mice with EMS

Peritoneal fluid was collected from 6-8-week-old female mice as described above. Cells were incubated with the F4/80 antibody on ice for 30 minutes, and macrophages were subsequently sorted using a Beckman Coulter flow cytometer. After establishing endometriosis in C57BL/6JGpt recipient mice, 1×10⁶ peritoneal macrophages pre-treated with either the STING agonist diABZI (5 µM) or PBS were administered intraperitoneally on days 10 and 20 post-induction. Estrogen (0.2 µg/g in 200µl PBS) was injected every three days, starting from day 4. All recipient mice were euthanized for analysis four weeks after model establishment.

### 10. Flow cytometry

Peritoneal lavage fluid was collected and filtered through a 70-µm strainer to obtain a single-cell suspension. Red blood cells were lysed, and approximately 1×10⁶ cells per sample were stained with surface antibodies (see Supplementary Materials) at optimized concentrations on ice for 15-20 minutes in the dark. After two washes, cells were fixed and permeabilized using an intracellular fixation and permeabilization buffer (ThermoFisher) and incubated with intracellular antibodies on ice for 30 minutes. Data were acquired on a BD LSRFortessa flow cytometer and analyzed using FlowJo software (BD Biosciences).

### 11. H&E

Hematoxylin and eosin (H&E) staining was performed according to a standard protocol^63^.

### 12. Multi-color immunohistochemistry

Multiplex fluorescent immunohistochemistry was carried out using a four-color staining kit (Absin, abs50028) following the manufacturer’s instructions. After dewaxing and hydration, paraffin sections of endometriotic lesions were fixed in 10% neutral formalin for 10 minutes, followed by antigen retrieval via microwave heating in retrieval solution for 15 minutes. Sections were blocked with 5% serum for 10 minutes and incubated with primary antibodies for 1 hour at room temperature.

After TBST washes, sections were incubated with HRP-conjugated secondary antibodies for 10 minutes, followed by TSA (1:100) and DAPI staining, each for 10 minutes at room temperature. Images were captured using a confocal microscope (Nikon) and NIS-Elements Viewer software and analyzed with ImageJ. Antibody details are provided in the Supplementary Materials.

### 13. Isolation of bone marrow-derived macrophages

Femurs and tibias were isolated from 6-10 week-old mice, and bone marrow cells were flushed using a 5-ml syringe. The cell suspension was filtered through a 70-µm strainer and centrifuged at 600g for 5 minutes. Red blood cells were lysed using a lysis buffer. Cells were cultured in RPMI-1640 medium containing 10% FBS, 1% penicillin-streptomycin, and 10 ng/ml M-CSF. Medium was replenished on days 3 and 5. On day 6, bone marrow-derived macrophages (BMDMs) were used for experiments.

### 14. Reagents and ELISA

Peritoneal macrophages and BMDMs were treated with 5 µg/ml digitonin for 12 hours, followed by stimulation with 10 µg/ml 2’3’-cGAMP for 8 hours. Supernatants were collected, and IFN-β levels were measured using a Mouse IFN-β ELISA Kit (4A BIOTECH, CME0116) according to the manufacturer’s instructions.

### 15. Immunoblotting and coimmunoprecipitation

For immunoblotting, cells were lysed on ice for 30 minutes using RIPA buffer (Thermo Scientific, 89900) supplemented with protease and phosphatase inhibitors (Thermo Scientific, 1861281). Proteins were separated by SDS-PAGE and transferred to PVDF membranes. Membranes were blocked with 5% skim milk in TBST for 1 hour at room temperature and incubated with primary antibodies overnight at 4°C. After washing, membranes were incubated with HRP-conjugated secondary antibodies for 1 hour at room temperature and visualized using a chemiluminescence imager (Tanon).

For co-immunoprecipitation, cells were lysed in IP lysis buffer (Thermo Scientific, 87787) with inhibitors. Lysates were incubated with primary antibodies overnight at 4°C with rotation. Beads were blocked with 5% BSA (ABCONE, 9048-46-8) for 1 hour and incubated with antigen-antibody complexes for 6 hours at 4°C. Proteins were eluted under denaturing conditions and analyzed by immunoblotting.

### 16. Confocal microscopy

Peritoneal macrophages were cultured on sterile coverslips in 12-or 24-well plates. After treatment, cells were fixed with 4% PFA for 20 minutes, permeabilized with 0.5% Triton X-100 for 10 minutes, and blocked with 5% goat serum for 1 hour. Primary antibody incubation was performed for 2 hours at 37°C or overnight at 4°C. Cells were then incubated with HRP-conjugated secondary antibodies for 30 minutes, followed by TSA reaction (1:100) for 10 minutes. Antibodies were eluted with preheated elution buffer (37°C) for 30 minutes. After re-blocking, the procedure was repeated for subsequent antibodies. Coverslips were stained with DAPI and mounted for imaging. Images were acquired using a confocal microscope.

### 17. Human samples

Endometriotic and healthy control tissues were obtained from informed consenting donors. The study was approved by the Institutional Research Ethics Committee of Southern Medical University (approval number: 2024-KY-273-01).

### 18. Statistical analysis

Data were analyzed using GraphPad Prism 9.0, SPSS 24.0, or R software. Comparisons between two groups were made using unpaired Student’s t-tests; one-way ANOVA was used for multiple groups. A p-value < 0.05 was considered statistically significant. Data are presented as mean ± SEM or SD from at least three independent experiments.

## Acknowledgments

This study was supported by the grants from the National Natural Science Foundation of China (82073165), Natural Science Foundation of Guangdong province (2214050008966, 20257625412,26842516959), the Medical Science and Technology Research Foundation of Guangdong Province(A2023124).

## Author’s Contributions

S Wu, W Zhu and LW Hao designed the research, QL Mo, RS Chang and LY Zhang performed experiments and wrote the manuscript. W Huang, QB Zhang, XY Liang, XL Xue, XR Hou, YC Lin, ZL Zhou and YW Chen assisted in performing experiments. L Huang provided suggestions.

## Data Availability

The raw data supporting the conclusions of this article will be made available by the authors, without undue reservation, to any qualified researcher.

## References

1. Chapron C, Marcellin L, Borghese B, Santulli P. Rethinking mechanisms, diagnosis and management of endometriosis. Nat Rev Endocrinol. 2019;15(11):666–682

2. Zondervan KT, Becker CM, Koga K et al. Endometriosis. Nat Rev Dis Primers. 2018;4(1):9

3. Kvaskoff M, Mahamat-Saleh Y, Farland LV et al. Endometriosis and cancer: a systematic review and meta-analysis. Hum Reprod Update. 2021;27(2):393–420

4. Nezhat C, Falik R, McKinney S, King LP. Pathophysiology and management of urinary tract endometriosis. Nat Rev Urol. 2017;14(6):359–372

5. He ZX, Shi HH, Fan QB et al. Predictive factors of ovarian carcinoma for women with ovarian endometrioma aged 45 years and older in China. J Ovarian Res. 2017;10(1):45

6. Hermens M, van Altena AM, Nieboer TE et al. Incidence of endometrioid and clear-cell ovarian cancer in histological proven endometriosis: the ENOCA population-based cohort study. Am J Obstet Gynecol. 2020;223(1):107.e1–107.e11

7. Taylor HS, Kotlyar AM, Flores VA. Endometriosis is a chronic systemic disease: clinical challenges and novel innovations. Lancet. 2021;397(10276):839–852

8. Ardavín C, Alvarez-Ladrón N, Ferriz M, Gutiérrez-González A, Vega-Pérez A. Mouse Tissue-Resident Peritoneal Macrophages in Homeostasis, Repair, Infection, and Tumor Metastasis. Adv Sci (Weinh). 2023;10(11):e2206617

9. Lei ST, Lai ZZ, Hou SH et al. Abnormal HCK/glutamine/autophagy axis promotes endometriosis development by impairing macrophage phagocytosis. Cell Prolif. 2024;57(11):e13702

10. Huang YL, Zhang FL, Tang XL, Yang XJ. Telocytes Enhances M1 Differentiation and Phagocytosis While Inhibits Mitochondria-Mediated Apoptosis Via Activation of NF-κB in Macrophages. Cell Transplant. 2021;30:9636897211002762

11. Hou HT, Lin TC, Wu MH, Tsai SJ. Feel so bac: is Fusobacterium the suspect causing endometriosis? Trends Mol Med. 2023;29(10):780–782

12. Bai X, Guo YR, Zhao ZM et al. Macrophage polarization in cancer and beyond: from inflammatory signaling pathways to potential therapeutic strategies. Cancer Lett. 2025;625:217772

13. Yan L, Wang J, Cai X et al. Macrophage plasticity: signaling pathways, tissue repair, and regeneration. MedComm (2020). 2024;5(8):e658

14. Gou Y, Li X, Li P et al. Estrogen receptor beta upregulates CCL2 via NF-kappaB signaling in endometriotic stromal cells and recruits macrophages to promote the pathogenesis of endometriosis. Hum Reprod. 2019;34(4):646–658

15. Sun SG, Guo JJ, Qu XY et al. The extracellular vesicular pseudogene LGMNP1 induces M2-like macrophage polarization by upregulating LGMN and serves as a novel promising predictive biomarker for ovarian endometriosis recurrence. Hum Reprod. 2022;37(3):447–465

16. Li H, Chai X. PDPK1 governs macrophage M2 polarization via hypoxia-driven CD47/AKT-glycolytic Axis in endometriosis. Cell Signal. 2025;134:111922

17. Martinez-Zamora MA, Armengol-Badia O, Quintas-Marques L, Carmona F, Closa D. Macrophage Phenotype Induced by Circulating Small Extracellular Vesicles from Women with Endometriosis. Biomolecules. 2024;14(7)

18. Chen C, Xu P. Cellular functions of cGAS-STING signaling. Trends Cell Biol. 2023;33(8):630–648

19. Sun X, Liu L, Wang J et al. Targeting STING in dendritic cells alleviates psoriatic inflammation by suppressing IL-17A production. Cell Mol Immunol. 2024;21(7):738–751

20. Liu Y, Carmona-Rivera C, Seto NL et al. Role of STING Deficiency in Amelioration of Mouse Models of Lupus and Atherosclerosis. Arthritis Rheumatol. 2024;

21. Ni L, Lin Z, Hu S et al. Itaconate attenuates osteoarthritis by inhibiting STING/NF-κB axis in chondrocytes and promoting M2 polarization in macrophages. Biochem Pharmacol. 2022;198:114935

22. Geng K, Ma X, Jiang Z et al. High glucose-induced STING activation inhibits diabetic wound healing through promoting M1 polarization of macrophages. Cell Death Discov. 2023;9(1):136

23. Wang Q, Bergholz JS, Ding L et al. STING agonism reprograms tumor-associated macrophages and overcomes resistance to PARP inhibition in BRCA1-deficient models of breast cancer. Nat Commun. 2022;13(1):3022

24. Li Y, Yi J, Ma R et al. A polymeric nanoplatform enhances the cGAS-STING pathway in macrophages to potentiate phagocytosis for cancer immunotherapy. J Control Release. 2024;373:447–462

25. Bulun SE. Endometriosis. N Engl J Med. 2009;360(3):268–79

26. Aydin Y, Atis A, Ercan E, Donmez M. An endometriotic vault fistula presenting with monthly bleeding after hysterectomy. Arch Gynecol Obstet. 2009;280(6):1011–4

27. Burney RO, Lathi RB. Menstrual bleeding from an endometriotic lesion. Fertil Steril. 2009;91(5):1926–7

28. Sharma R, Antypiuk A, Vance SZ et al. Macrophage metabolic rewiring improves heme-suppressed efferocytosis and tissue damage in sickle cell disease. Blood. 2023;141(25):3091–3108

29. Tenhunen R, Marver HS, Schmid R. The enzymatic conversion of heme to bilirubin by microsomal heme oxygenase. Proc Natl Acad Sci U S A. 1968;61(2):748–55

30. Campbell NK, Fitzgerald HK, Dunne A. Regulation of inflammation by the antioxidant haem oxygenase 1. Nat Rev Immunol. 2021;21(7):411–425

31. Imanaka S, Yamada Y, Kawahara N, Kobayashi H. A delicate redox balance between iron and heme oxygenase-1 as an essential biological feature of endometriosis. Arch Med Res. 2021;52(6):641–647

32. Ma XQ, Liu YY, Zhong ZQ et al. Heme induced progesterone-resistant profiling and promotion of endometriosis in vitro and in vivo. Biochim Biophys Acta Mol Basis Dis. 2023;1869(7):166761

33. Van Langendonckt A, Casanas-Roux F, Dolmans MM, Donnez J. Potential involvement of hemoglobin and heme in the pathogenesis of peritoneal endometriosis. Fertil Steril. 2002;77(3):561–70

34. Ogawa K, Liu T, Kawahara N, Kobayashi H. Macrophages Protect Endometriotic Cells Against Oxidative Damage Through a Cross-Talk Mechanism. Reprod Sci. 2022;29(8):2165–2178

35. Lu JJ, Ning Y, Hu WT et al. Excess heme orchestrates progesterone resistance in uterine endometrial cancer through macrophage polarization and the IL-33/PAX8/PGR axis. Biomed Pharmacother. 2025;186:118008

36. Sha W, Zhao B, Wei H et al. Astragalus polysaccharide ameliorates vascular endothelial dysfunction by stimulating macrophage M2 polarization via potentiating Nrf2/HO-1 signaling pathway. Phytomedicine. 2023;112:154667

37. Chen W, Zhang Y, Chen J et al. Heme Oxygenase-1 Modulates Macrophage Polarization Through Endothelial Exosomal miR-184-3p and Reduces Sepsis-Induce Lung Injury. Int J Nanomedicine. 2025;20:5039–5057

38. Wu Y, Wu B, Zhang Z et al. Heme protects intestinal mucosal barrier in DSS-induced colitis through regulating macrophage polarization in both HO-1-dependent and HO-1-independent way. FASEB J. 2020;34(6):8028–8043

39. Bulun SE, Yilmaz BD, Sison C et al. Endometriosis. Endocr Rev. 2019;40(4):1048–1079

40. Liu ZZ, Zhou CK, Lin XQ et al. STING-dependent trained immunity contributes to host defense against Clostridium perfringens infection via mTOR signaling. Vet Res. 2024;55(1):52

41. Lee SJ, Yang H, Kim WR et al. STING activation normalizes the intraperitoneal vascular-immune microenvironment and suppresses peritoneal carcinomatosis of colon cancer. J Immunother Cancer. 2021;9(6)

42. Yuan M, Li D, An M et al. Rediscovering peritoneal macrophages in a murine endometriosis model. Hum Reprod. 2017;32(1):94–102

43. Su W, Gao W, Zhang R et al. TAK1 deficiency promotes liver injury and tumorigenesis via ferroptosis and macrophage cGAS-STING signalling. JHEP Rep. 2023;5(5):100695

44. Moore JA, Mistry JJ, Hellmich C et al. LC3-associated phagocytosis in bone marrow macrophages suppresses acute myeloid leukemia progression through STING activation. J Clin Invest. 2022;132(5)

45. Zhu S, Chen Q, Sun J et al. The cGAS-STING pathway promotes endometriosis by up-regulating autophagy. Int Immunopharmacol. 2023;117:109644

46. Xu Z, Zhao H, Yue C et al. Low STING expression promotes endometrial stromal cell invasion and migration via the STING/IRF-3/IFN-β1 pathway in eutopic endometrium of women with endometriosis. Gynecol Endocrinol. 2022;38(12):1129–1135

47. Kwon J, Bakhoum SF. The Cytosolic DNA-Sensing cGAS-STING Pathway in Cancer. Cancer Discov. 2020;10(1):26–39

48. Motwani M, Pesiridis S, Fitzgerald KA. DNA sensing by the cGAS-STING pathway in health and disease. Nat Rev Genet. 2019;20(11):657–674

49. Zhang C, Deng Z, Wu J et al. HO-1 impairs the efficacy of radiotherapy by redistributing cGAS and STING in tumors. J Clin Invest. 2024;134(23)

50. Sosnowska D, Cheung TS, Sarkar J et al. Tumor microenvironments with an active type-I interferon response are sensitive to inhibitors of heme degradation. JCI Insight. 2025;

51. Pais TF, Ali H, Moreira DSJ et al. Brain endothelial STING1 activation by Plasmodium-sequestered heme promotes cerebral malaria via type I IFN response. Proc Natl Acad Sci U S A. 2022;119(36):e2206327119

52. Hogg C, Panir K, Dhami P et al. Macrophages inhibit and enhance endometriosis depending on their origin. Proc Natl Acad Sci U S A. 2021;118(6)

53. Horne AW, Missmer SA. Pathophysiology, diagnosis, and management of endometriosis. BMJ. 2022;379:e070750

54. Stuart T, Butler A, Hoffman P et al. Comprehensive Integration of Single-Cell Data. Cell. 2019;177(7):1888–1902.e21

55. Korsunsky I, Millard N, Fan J et al. Fast, sensitive and accurate integration of single-cell data with Harmony. Nat Methods. 2019;16(12):1289–1296

56. Zou G, Wang J, Xu X et al. Cell subtypes and immune dysfunction in peritoneal fluid of endometriosis revealed by single-cell RNA-sequencing. Cell Biosci. 2021;11(1):98

57. Ma J, Zhang L, Zhan H et al. Single-cell transcriptomic analysis of endometriosis provides insights into fibroblast fates and immune cell heterogeneity. Cell Biosci. 2021;11(1):125

58. Hao Y, Stuart T, Kowalski MH et al. Dictionary learning for integrative, multimodal and scalable single-cell analysis. Nat Biotechnol. 2024;42(2):293–304

59. Gulati GS, Sikandar SS, Wesche DJ et al. Single-cell transcriptional diversity is a hallmark of developmental potential. Science. 2020;367(6476):405-411

60. Weiler P, Lange M, Klein M, Pe’Er D, Theis F. CellRank 2: unified fate mapping in multiview single-cell data. Nat Methods. 2024;21(7):1196–1205

61. Trapnell C, Cacchiarelli D, Grimsby J et al. The dynamics and regulators of cell fate decisions are revealed by pseudotemporal ordering of single cells. Nat Biotechnol. 2014;32(4):381–386

62. Ray A, Dittel BN. Isolation of mouse peritoneal cavity cells. J Vis Exp. 2010;(35)

63. Yang XY, Yu H, Fu J et al. Hydroxyurea ameliorates atherosclerosis in ApoE(−/−) mice by potentially modulating Niemann-Pick C1-like 1 protein through the gut microbiota. Theranostics. 2022;12(18):7775–7787

