## Supplementary files for "Hemin Inhibits the Activation of STING in Macrophages by Inducing HO-1, Promoting Endometriosis Development"

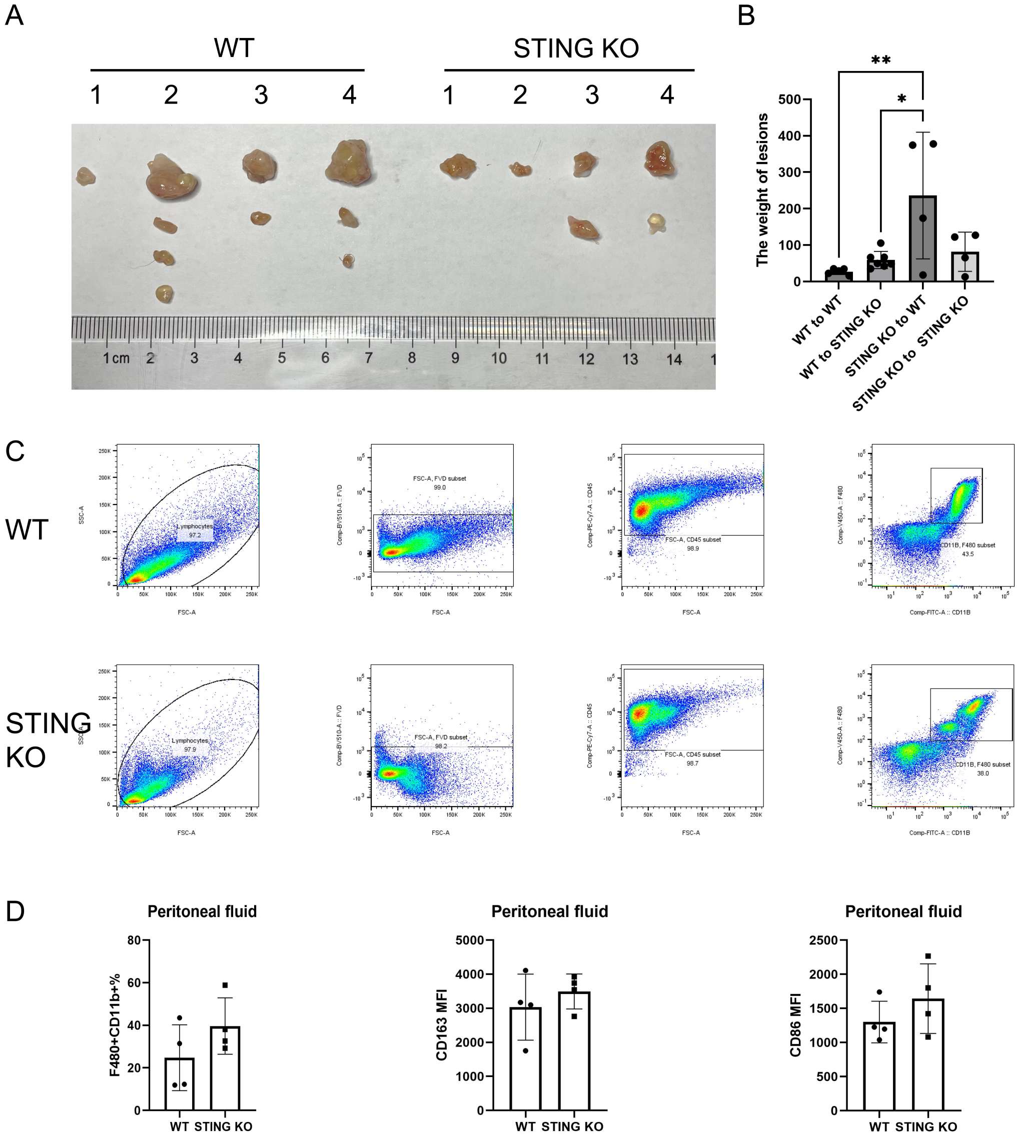


Supplementary figure1

A Macroscopic aspect of lesions at 1 month from mice with induced EMS using STING$^{-/-}$ mice as donors.

B The quantification of celll number and viability in lesions from WT mice or STING$^{-/-}$ mice.

C Flow plot indicating gating of macrophages (single/live/CD45$^{+}$F4/80$^{+}$CD11b$^{+}$) in peritoneal macrophages of mice with induced EMS.

D The quantification of F4/80$^{+}$CD11b$^{+}$ macrophage and the mean fluorescence intensity (MFI) of CD86/CD163 of F4/80$^{+}$CD11b$^{+}$ macrophage.

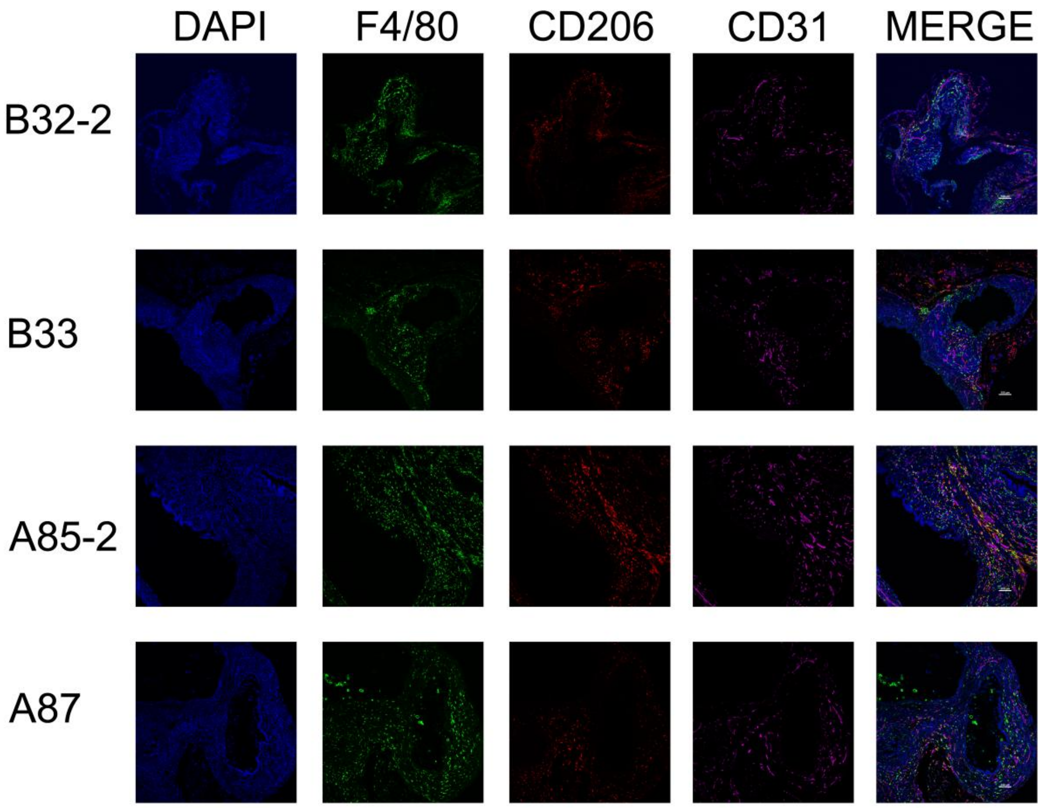


Supplementary figure2

Representative mIHC of EMS lesions. DAPI(blue), F4/80 (green), CD206 (red), CD31 (magenta). Bar = 100 μm. B32/B33 were from WT mice with induced EMS. A85/A87 were from STING$^{-/-}$ mice with induced EMS.

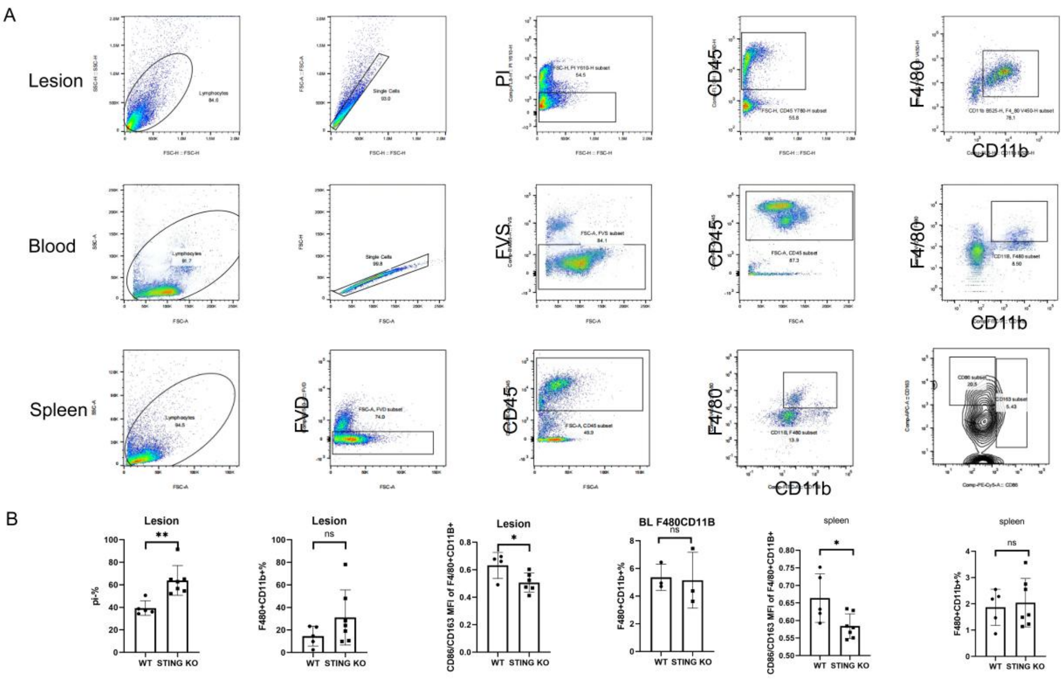


Supplementary figure3

A Flow plot indicating gating of macrophages (single/live/CD45$^{+}$/F4/80$^{+}$CD11b$^{+}$) in EMS lesions, peripheral blood and spleen respectively from STING$^{-/-}$ mice or WT mice with induced EMS.

B The quantification of F4/80$^{+}$CD11b$^{+}$ macrophage and mean fluorescence intensity(MFI) of CD86/CD163. *P＜0.05

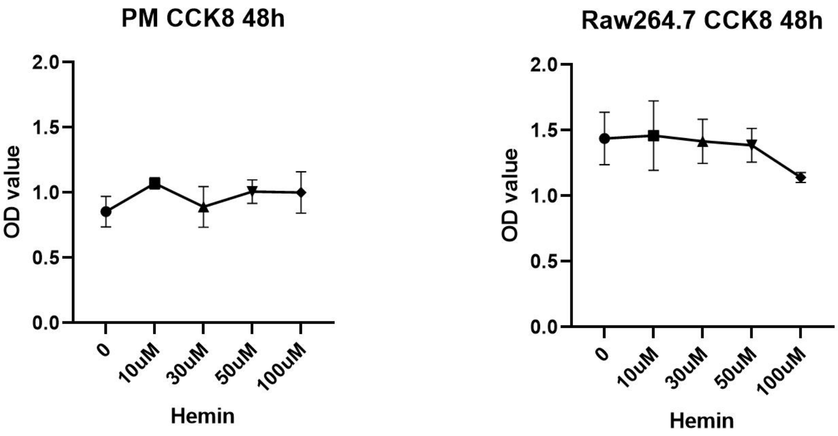


Supplementary figure 4

CCK8 assay of peritoneal macrophages and Raw264.7 treated with hemin after 48 hours.

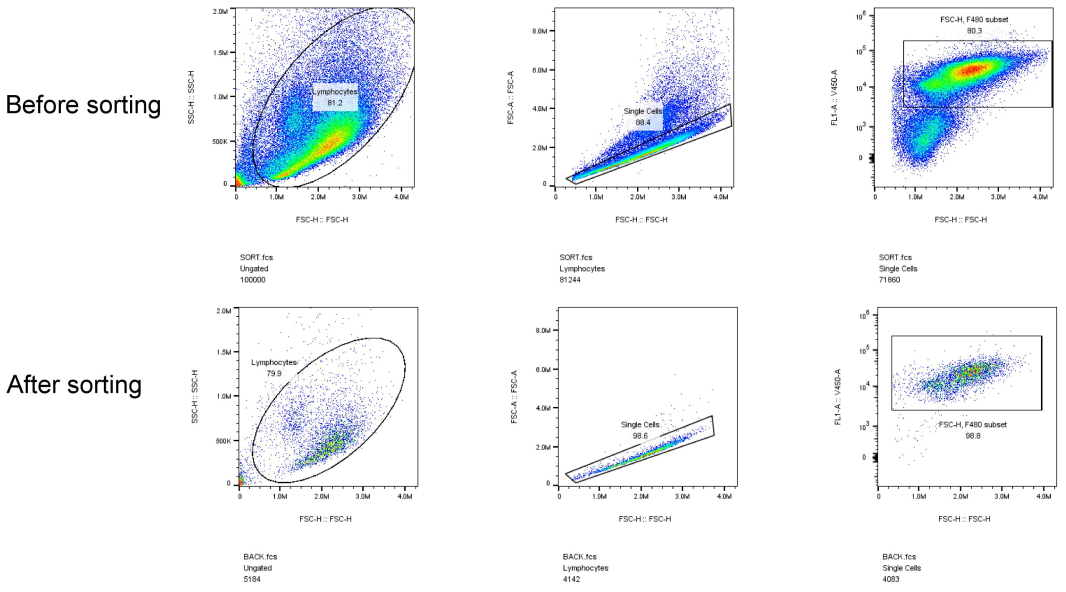


Supplementary figure5

Macrophages were subsequently sorted using a Beckman Coulter flow cytometer.

A

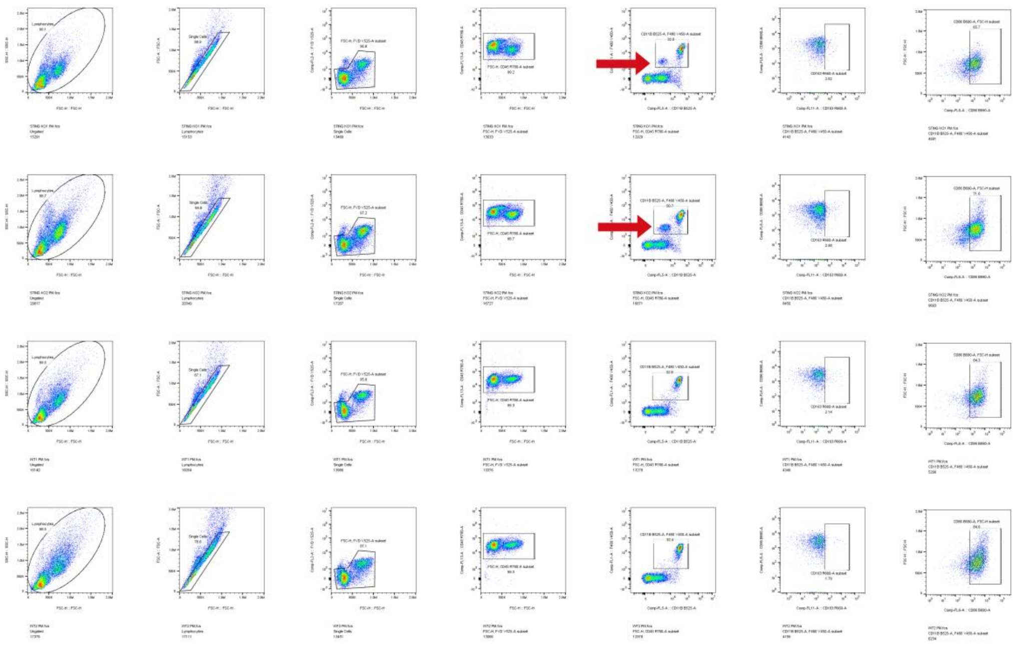

B

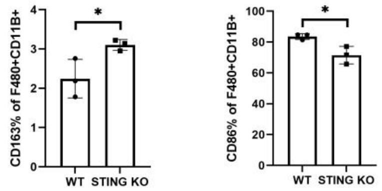


C

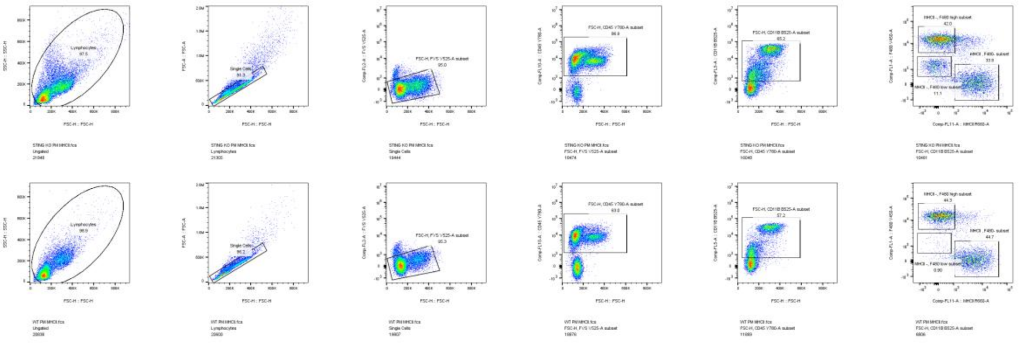

Supplementary figure6

A Flow plot of peritoneal fluid of mice from untreated WT mice or STING$^{-/-}$ mice.

B Unprocessed statistical graph of the percentage CD163$^{+}$ in F4/80$^{+}$CD11b$^{+}$ macrophages.

C Unprocessed flow plot of peritoneal fluid of mice from WT mice or STING$^{-/-}$ mice.
